# Histone deacetylase inhibitors correct the cholesterol storage defect in most NPC1 mutant cells

**DOI:** 10.1101/076695

**Authors:** Nina H. Pipalia, Kanagaraj Subramanian, Shu Mao, William E. Balch, Frederick R. Maxfield

**Affiliations:** Department of Biochemistry, Weill Cornell Medical College, New York, NY 10065, USA; Department of Chemical Physiology, and Cell and Molecular Biology. The Skaggs Institute for Chemical Biology, The Scripps Research Institute, La Jolla, California, 92037, USA

**Author notes:** These authors contributed equally to this work (Nina H. Pipalia, Kanagaraj Subramanian, Shu Mao). Frederick R. Maxfield Department of Biochemistry Weill Cornell Medical College 1300 York Avenue New York, NY 10065. William E. Balch Department of Chemical Physiology, and Cell and Molecular Biology The Skaggs Institute for Chemical Biology The Scripps Research Institute La Jolla, California, 92037.

**Keywords:** Niemann Pick type C, Cellular cholesterol, lipid transport, Inborn errors of metabolism, Drug therapy, Cholesterol trafficking

## Abstract

Niemann Pick C disease (NPC) is an autosomal recessive disorder that leads to excessive storage of cholesterol and other lipids in late endosomes and lysosomes. The large majority of NPC disease is caused by mutations in NPC1, a large polytopic membrane protein that functions in late endosomes. There are many disease-associated mutations in NPC1, and most patients are compound heterozygotes. The most common mutation NPC1^I1061T^ has been shown to cause endoplasmic reticulum associated degradation of the NPC1 protein. Treatment of patient derived NPC1^I1061T^ fibroblasts with histone deacetylase inhibitors (HDACi) Vorinostat or Panobinostat increases expression of the mutant NPC1 protein and leads to correction of the cholesterol storage. Herein we show that several other human NPC1 mutant fibroblast cell lines can also be corrected by Vorinostat or Panobinostat and that treatment with Vorinostat extends the lifetime of the NPC1^I1061T^ protein. To test effects of HDACi on a large number of *NPC1* mutants, we engineered a U2OS cell line to suppress NPC1 expression by shRNA and then transiently transfected these cells with 81 different NPC1 mutant constructs. The mutant NPC1 did not significantly reduce cholesterol accumulation, but approximately 80% of the mutants showed reduced cholesterol accumulation when treated with Vorinostat or Panobinostat.

## Introduction

Lipoprotein-derived cholesterol is normally transported out of late endosomes and lysosome (LE/Ly) by a process that requires the NPC1 and NPC2 proteins (1, 2). A current model (3) suggests that cholesterol, which is a product of the hydrolysis of cholesteryl esters by lysosomal acid lipase, is first transferred to NPC2, a cholesterol binding protein. The cholesterol can then be transferred to the N-terminal domain of NPC1, a LE/Ly membrane protein with 13 transmembrane segments (4). This model is supported by recent structural studies of NPC1 (5, 6) and of NPC2 bound to the middle lumenal domain of NPC1 (7). Cholesterol finally leaves the LE/Ly by a process that is not well understood. In cells with defects in NPC1 or NPC2, cholesterol and other lipids accumulate in the LE/Ly. Cells with these defects show multiple defects in lipid and protein trafficking (8, 9).

Defects in NPC1 or NPC2 in humans lead to an autosomal recessive lysosomal storage disease, Niemann-Pick C (NPC) disease, which causes pathology in multiple tissues but especially in the central nervous system (10, 11). A high fraction of patients with NPC disease die of neurological complications before the age of 25 (10). In addition to cholesterol other lipids, including glycosphingolipids, accumulate in the LE/Ly of patient cells. There are currently no effective treatments for NPC disease approved by the U.S. Food and Drug Administration (FDA). Miglustat, an inhibitor of glycosphingolipid synthesis (12), shows some beneficial effects and is approved for use in several countries (13, 14). Treatment of mice and cats with hydroxypropyl-beta-cyclodextrin (HPBCD) has been shown to effectively reduce the storage of cholesterol and other lipids in cells and to ameliorate symptoms (15–17). HPBCD does not cross the blood brain barrier, so in cats and humans it needs to be injected directly into the CNS. Early stage clinical trials of HPBCD have been carried out in humans (18).

In a previous study (19) we showed that HDACi including Vorinostat (also called Suberoylanilide hydroxamic acid or SAHA) and Panobinostat (LBH589) are remarkably effective in correcting the NPC1 phenotype in human fibroblast cells that have an *NPC1*^*I1061T*^ mutation. The pharmacological profile was most consistent with the effects being attributed to inhibition of HDACs 1, 2, or 3 (20). Treatment of patient-derived fibroblasts with HDACi reduced the accumulation of cholesterol in lysosomal storage organelles (LSOs) and restored other aspects of cholesterol homeostasis including normal processing of sterol regulatory element-binding protein 2 (SREBP2) and reduction of the expression of LDL receptors (19, 21). HDACi treatment did not correct the cholesterol storage defect of patient-derived cells expressing *NPC2* mutations (19), indicating that the HDACi do not bypass the need for the NPC1/NPC2 transport system as HPBCD does (22). This indicated that the HDACi might work by allowing the mutant NPC1 proteins to function sufficiently well to correct the cholesterol transport out of LSOs.

The mechanism by which HDACi might restore the function of mutant NPC1 proteins has not been determined. It has been observed that there is more rapid degradation of the NPC1^I1061T^ protein as compared to wild type (WT) NPC1 protein, and it was proposed that this is due to enhanced endoplasmic reticulum associated degradation (ERAD) of the mutant protein (23). Treatment of cells expressing NPC1^I1061T^ with HDACi such as Panobinostat or Vorinostat increased the expression of the mutant NPC1 protein (19). Correction of the NPC phenotype would require that this mutant protein retains adequate functional capability and that a sufficient amount is delivered to the LE/Ly. Other data are consistent with the hypothesis that some mutant NPC1 proteins can function in LE/Ly if they are delivered to those organelles. Simply overexpressing NPC1^I1061T^ in mutant cells leads to partial correction of the phenotype (23). Some indirect treatments also increase the abundance of NPC1 and lead to correction of the phenotype in cultured cells. These include treatment with ryanodine receptor antagonists (24), treatment with oxysterols that bind to NPC1 (25), or reduced expression of TMEM97, an NPC1-binding protein (26). These studies have indicated that alterations in the proteostasis environment by various mechanisms leads to reduced degradation of mutant forms of NPC1. As described herein, we found that treatment of some NPC1 mutant cells with Vorinostat led to a longer lifetime of the NPC1^I1061T^ protein and increased delivery of the protein to LE/Ly. A recent study in mice reported that a combination therapy with Vorinostat and HPBCD led to slowed neuronal degeneration and improved lifespan in Npc1 mutant animals (27).

Approximately 95% of NPC cases are due to mutations in the NPC1 protein, and the *NPC1*^*I1061T*^ mutation, which occurs in about 15-20% of NPC1 patients, is the most commonly observed mutation (28, 29). However, more than 300 different *NPC1* mutations have been observed that are known to be or are likely to be pathogenic (10, 30). It would be very difficult to test drug treatments in hundreds of different human NPC1 mutant fibroblast cell lines, and the large number of compound heterozygous mutations would make it nearly impossible to evaluate the ability of HDACi to correct a specific mutation. In order to evaluate the effectiveness of HDACi as a potential therapy for NPC patients we developed an efficient screening system using an engineered cell line. Human U2OS osteosarcoma cells were stably transfected with Scavenger Receptor type A (SRA), and the endogenous NPC1 expression in the cells was stably silenced with an shRNA. The U2OS-SRA-shNPC1 cells were then transiently transfected with a bicistronic vector expressing GFP (to identify transfected cells) and one of 81 *NPC1* mutations found in patients. This system was used to test the effect of HDACi treatments on multiple NPC1 mutations simultaneously. After treatment with Vorinostat or Panobinostat, a high fraction of NPC1 mutant proteins were effective in reducing cholesterol accumulation. This suggests that HDACi therapy might be effective for a large majority of NPC1 patients. Since Vorinostat and Panobinostat are FDA-approved drugs for treatment of some cancers, and other HDACi have been in large scale clinical trials, HDACi may be considered as a potential therapy for NPC1 disease (20).

## Materials and Methods

### Reagents

Gibco^®^ McCoy's 5A Medium, Modified Eagle’s Medium (MEM), Fetal Bovine Serum (FBS), Hank’s Balanced Salt Solution (HBSS), penicillin/streptomycin (P/S), Geneticin (G418), AlexaFluor-546, Cy5 Goat anti-Rat (A10525), and Alexa Fluor 546 Goat anti-Rabbit and Alexa Flour 488 Goat anti-Rat antibody, 1.1’-Dioctadecyl-3,3,3’3’-Tetramethyindocarbocyanine perchlorate (DiIC18(3)) were purchased from Invitrogen Life Technologies Corporation (Carlsbad, CA). Effectene Transfection Reagent Kit and DNA purification kit was purchased from QIAGEN Inc. (Valencia, CA). HDACi’s (Vorinostat and Panobinostat) were stocked at 5 mM in Dimethyl Sulfoxide (DMSO) and stored at −20°C. Vorinostat and Panobinostat were a generous gift from Dr. Paul Helquist (University of Notre Dame, South Bend, IN). Acetylated low-density lipoprotein (AcLDL) was prepared by acetylation of low-density lipoprotein with acetic anhydride (31). AlexaFluor-546 labeled human low-density lipoprotein (LDL-Alexa546) was prepared as described (32, 33). Rabbit polyclonal anti-NPC1 antibody and rabbit polyclonal anti-LAMP1 antibody (ab24170) were purchased from Abcam (Cambridge, MA). All other chemicals, including (DMSO, 99% fatty-acid free bovine serum albumin (BSA), filipin, paraformaldehyde (PFA) and 4-(2-hydroxyethyl)-1-piperazineethanesulfonic acid (HEPES) were purchased from Sigma Chemical (St. Louis, MO). Draq5 was from Biostatus (Leicestershire, UK), Metamorph image-analysis software was from Molecular Devices (Downington, PA).

### Human fibroblast cells

Human NPC1 fibroblasts GM05659, GM18453 (homozygous *NPC1* mutant *I1061T*), and GM03123 (heterozygous *NPC1* mutations *P237S* and *I1061T)* were from Coriell Institute, (Camden, NJ). (The P237S allele has been found recently to be in linkage with a pathogenic splice mutation that may be responsible for its defect (F.D. Porter, submitted).) Patient derived mutant NPC1 skin fibroblasts were from Forbes Porter’s laboratory at the National Institutes of Health (Bethesda, MD) as listed in Table 1. All human skin fibroblasts were maintained in Modified Eagle’s Medium (MEM) supplemented with 10% FBS. For drug treatment cells were maintained in MEM supplemented with 5.5% FBS and 20 mM HEPES.

### U2OS cells stably expressing of SRA (U2OS-SRA)

Human osteosarcoma U2OS (HTB-96^™^) cells from ATCC were transfected with murine scavenger receptor type IIA (SRA-II) cDNA [Kind gift from Dr. Monty Krieger (Massachusetts Institute of Technology, Boston, MA)]. Briefly, the 1.0 kb coding region of murine scavenger receptor type II was cut from a larger 4.0 kb plasmid encoding the coding region and the 3’ untranslated region in a pcDNA3.1 expression vector (34). The coding sequence was amplified using polymerase chain reaction (PCR) and inserted into a neomycin resistant pcDNA 3.1 TOPO plasmid using pcDNA 3.1 Directional TOPO Expression kit (Invitrogen, Carlsbad, CA) according to the manufacturer’s instructions. The insertion of SRA-II in the pcDNA3.1 TOPO plasmid was confirmed by band size of full length plasmid and also after digestion with appropriate restriction enzymes using gel electrophoresis. The pcDNA 3.1 TOPO plasmid encoding SRA-II was purified using Qiagen DNA purification kit (Qiagen Inc. Valencia, CA) and transfected in U2OS cells using Lipofectamine (Invitrogen, Carlsbad, CA) as a transfection reagent. U2OS cells expressing murine SRA-II were grown in McCoy's 5A medium supplemented with 10% FBS, 1% P/S and selection antibiotic G418 (1 mg/ml) in a humidified incubator with 5% CO_2_ at 37°C for five passages. To select a population of U2OS cells expressing the SRA-II gene, cells were incubated with DiIC18(3) labeled AcLDL and sorted using flow cytometry cell sorting (FACS). Murine macrophage cells J774 were used as a positive control, and the pool of cells with equivalent DiI intensity, indicative of SRA-II expression, were collected. The cells were expanded, and stock cultures were frozen for future experiments.

### Effect of HDAC inhibitor treatment on human fibroblasts

The dose dependence of two HDACis (Vorinostat and Panobinostat) was determined after 48 hours treatment of NPC1 fibroblasts from several patients carrying different mutations. Human fibroblasts were seeded in - 384 well plates at 450 cells/ well in growth medium on day 1. Four cell lines were seeded in different wells of a plate, and GM03123 fibroblasts were used as a control in each plate. After overnight incubation, 2X concentrated compounds were added at six different doses such that the final concentration ranged from 40 nM to 10 μM for Vorinostat and 5 nM to 1 μM for Panobinostat diluted in growth medium supplemented with 20 mM HEPES buffer and 5.5% FBS. DMSO was used as a control in each plate for each concentration. After 48 hours, the plate was washed with PBS three times, fixed with 1.5% PFA, and stained with 50 μg/ml filipin and nuclear stain Draq5. Measurements were made from 4 wells for each condition in each experiment, and the experiment was repeated three times. Images were acquired on an ImageXpress^Micro^ automatic fluorescence microscope, at four sites per well and analyzed to obtain the LSO compartment ratio, which is a measure of filipin labeling of stored cholesterol (35). The LSO compartment ratio for each concentration was normalized to corresponding DMSO treated control.

### Persistence of Vorinostat effects

NPC1 human fibroblasts GM03123 and GM18453 were seeded in four different 384 well plates at 450 cells/ well in growth medium on day 1. After overnight incubation, compounds diluted in growth medium supplemented with 20 mM HEPES buffer and 5.5% FBS were added in three of the four plates to achieve the desired final concentrations. DMSO was used as a control in each plate for each concentration. After 3 days, Vorinostat-supplemented medium was aspirated from each plate and replaced with normal growth medium, and the cells were incubated for additional 0, 1, 2 or 3 days. At the end of each time point cells were stained with filipin, and the LSO ratio was determined. The experiment was performed twice independently. The LSO ratio for each concentration was normalized to corresponding DMSO treated value.

### Stable silencing of NPC1

A line of NPC1-deficient stable U2OS-SRA cells was generated by silencing the endogenous NPC1 expression with 3’-UTR shNPC1 lentivirus. A Mission shRNA clone was purchased from Sigma against 3’UTR (TRCN000000542) of human NPC1 (pLKO-shNPC1) for knockdown of NPC1 expression. Stable clones of NPC1-deficient cells were selected with 5 μg/ml puromycin for three weeks. Stable U2OS-SRA-shNPC1 cells were cultured in McCoy’s 5A medium supplemented with 10% FBS, 50-units/ml penicillin, 50 μg/ml streptomycin, 5 μg/ml puromycin and 1 mg/ml G418.

### NPC1 expression vector

cDNA encoding human ΔU3m*npc1*-WT construct was kindly provided by Dan Ory (Washington University, St Louis). The *NPC1* gene was subcloned into bicistronic retroviral plasmid, pMIEG3 using EcoR1 and Not1 restriction enzyme sites. pMIEG3 vector was generated from pMSCVneo vector (Clontech) in which the murine phosphoglycerate kinase (PKG) promoter and the Neomycin resistance (Neo ^r^) genes were replaced by IRES (Internal Ribosome Entry Site) and EGFP (enhanced green fluorescent protein) genes. The coexpression of *eGFP* with the *NPC1* gene enables detection of cells expressing the human NPC1 protein by fluorescent microscopy. Using WT pMIEG3-hNPC1 plasmid as a template, human NPC1 mutants were generated with Quick-Change XL Site-directed Mutagenesis Kit (Stratagene, La Jolla, CA). The PMIEG3 plasmid harbors an ampicillin resistance gene as a selection marker. These plasmids were transiently expressed in U2OS-SRA-shNPC1 cells.

### Reverse transfection, drug treatment and cholesterol loading of U2OS-SRA-shNPC1 cells

Using Effectene Transfection Reagent Kit, an expression vector containing one of the *NPC1* mutants (0.4 µg DNA) was mixed with 3.2 µl enhancer in 100 µl of DNA-condensation buffer (EC) buffer for 5 min. They were then mixed with 4µl Effectene in 100 µl EC buffer, and 5µl of this mixture was added to each well of 384-well plates using a Perkin Elmer Mini-JANUS liquid dispenser (PerkinElmer, Waltham, MA). The plates were centrifuged at 16 G for 10 min. U2OS-SRA-shNPC1 cells were prepared at 1.67×10^5^ cells/ml in McCoy's 5A medium supplemented with 5% FBS, 1% P/S (medium A), and 5000 cells (30 µl of U2OS-SRA-shNPC1 cells in medium A) were plated in each well of 384-well plates using a Titertek Multidrop 384 model 832 liquid dispenser (Thermo Fisher Scientific Inc., Waltham, MA). Cells were grown with 30 µl medium A in a humidified incubator with 5% CO_2_ at 37°C for 24 hours. After 24 hours cells were treated with Vorinostat (10µM from a 5 mM stock in DMSO) or Panobinostat (50 nM from a 5 mM DMSO stock) for 48 hours. Control cells were treated with equivalent volumes of DMSO. The drugs in DMSO were diluted in medium A. To obtain final concentration of 10 μM for Vorinostat treatment, 30 μl of the premixed 20µM Vorinostat in medium A was dispensed to each well using a Thermo Multi-Drop liquid dispenser. To obtain final concentration of 50 nM for Panobinostat treatment, 30 μl of the premixed 100 nM Panobinostat in medium A was added into each well of the plates containing cells. After 48 hours of treatment, cells were fixed with 1.5% PFA in PBS and stained with 50 µg/ml filipin in PBS (35). In order to load cells with cholesterol, cells were treated with 50 µg/ml AcLDL in medium A for 2 hours before fixation. All reagent additions, buffer changes and labeling with filipin were carried out robotically using a Thermo Multi-Drop liquid dispenser and a Bio-Tek Elx405 plate washer (Bio-Tek Instruments, Inc., Winooski, VT).

### Fluorescence Microscopy

An automated ImageXpress^Micro^ imaging system from Molecular Devices equipped with a 300W Xenon-arc lamp from Perkin-Elmer, a Nikon 10X Plan Fluor 0.3 numerical aperture (NA) objective, and a Photometrics CoolSnapHQ camera (1,392 x 1,040 pixels) from Roper Scientific was used to acquire images. Filipin images were acquired using 377/50 nm excitation and 447/60 nm emission filters with a 415 dichroic long-pass filter. GFP images were acquired using 472/30 nm excitation and 520/35 nm emission filters with a 669 dichroic long-pass (DCLP) filter. Filter sets assembled in Nikon filter cubes were obtained from Semrock.

### NPC1 lifetime measurements

Wild type GM05659 and NPC1^I1061T^ mutant GM18453 skin fibroblasts were treated with DMSO or Vorinostat for 48 hours in growth medium supplemented with 5.5% FBS. After 48 hours, cells were switched to cysteine/ methionine free medium supplemented with 2% dialyzed FBS for 1 hour followed by 1 hour incubation with ^35^S Cys/Met (150 μCi). The cells were subsequently washed twice with HBSS and chased in regular growth medium for times between 0-72 hours. Cell lysates were prepared at each time point, after washing cells twice with HBSS and lysing in non-denaturing lysis buffer (1% TritonX-100, 50 mM tris-Cl, pH 7.4, 300 mM NaCl, 5 mM EDTA, 0.02% sodium azide, 10 mM iodoacetamide, 1 mM PMSF, 2 μg/ml leupeptin). Separately antibody conjugated beads were prepared combining 50% protein A – Sepaharose bead slurry in ice-cold PBS and polyclonal rabbit antibody to NPC1 (Abcam – 6 μg/ml) and incubating the suspension at 4°C for 16 hours. Beads were washed three times with non-denaturing lysis buffer and resuspended in non-denaturing lysis buffer supplemented with 0.1% BSA. Lysed cells were immunoprecipitated by incubating for 2 hours at 4°C while mixing on tube rotator. Beads were centrifuged and washed three times with ice cold wash buffer (0.1% w/v triton X-100, 50 mM Tris-Cl, pH 7.4, 300 mM NaCl, 5 mM EDTA, 0.2% sodium azide). Beads were finally resuspended in 4X SDS buffer and centrifuged. Immunoprecipitated samples were run on 4-12% Bis-Tris gels (Nupage NP0321, Life Technologies). Dried gels were exposed to a phosphor imager screen (Amersham Bioscience) overnight. The screen was scanned using a Typhoon Trio PhosphorImager (GE Healthcare), and the images were saved as tiff files. The integrated intensity of the bands at the molecular weight corresponding to NPC1 protein was measured using Metamorph image analysis software. Within each condition, the intensities of bands corresponding to NPC1 protein for each time point were normalized to the corresponding zero time.

### Imaging and analysis

The 384-well plates were imaged for GFP and filipin using a 10X 0.3 NA dry objective on an ImageXpress^Micro^ automatic fluorescence microscope. Each well was imaged at four sites. Images were analyzed using MetaXpress image-analysis software. First, all images were corrected for slightly inhomogeneous illumination as described previously (35). A background intensity value was set as the fifth percentile intensity of each image, and this intensity value was subtracted from each pixel in the image. At the plating density used in this study, all fields maintain at least 5% of the imaged area cell-free. To identify the cells expressing GFP, a threshold value was chosen for each experiment and was applied to GFP images. Objects containing at least 540 pixels above threshold were selected as transfected cells.

For the GFP-expressing cells, a method described previously (35) was used to quantify the cholesterol accumulation in lysosomal storage organelles. This is expressed as an LSO ratio, which is the ratio of filipin fluorescence intensity in the brightly labeled center of the cells divided by the area of the cells. The ratio is determined using two thresholds that are applied to the filipin images. A low threshold is set to include all areas occupied by cells. The outlines of cells using the low threshold are similar to cell outlines in transmitted light images. A higher threshold is then set to identify regions brightly stained with filipin in cells. The thresholds were chosen for each experiment. The LSO ratio of transfected cells on a per image basis was determined using the equation below:

LSO Compartment Ratio = (Total intensity above high thresholded filipin intensity in GFP positive cells)/ (Number of pixels above low thresholded filipin intensity in GFP positive cells)

Each NPC1 mutant was transfected in 12 wells of a 384-plate. Six wells were treated with drug 10µM Vorinostat or 50 nM Panobinostat, and the other six were treated with DMSO as solvent control. WT and I1061T NPC1 were used as controls in each plate. There were three parallel plates for each experiment, and three independent experiments for each set of NPC1 mutants. The LSO ratio values for each experiment were normalized to the corresponding value for DMSO-treated I1061T NPC1 transfected cells. Normalized LSO ratio values from three independent experiments were averaged. The LSO ratio value for each NPC1 mutant with each treatment condition was averaged per image from 216 images (four sites × six wells × three plates × three experiments). In 216 images, there were about 40 to 120 images with GFP positive cells. On average there were three GFP positive cells per image, so 120 to 360 NPC1 expressing cells were averaged to get the LSO ratio value for each NPC1 mutant with each treatment condition. A standard error of mean (SEM) was calculated for each LSO ratio value. A p-value comparing LSO ratio values in DMSO treated cells and drug treated cells was calculated using t-test with two-tailed distribution and two-sample equal variance (homoscedastic) type using Microsoft Excel software.

### NPC1 immunolocalization in NPC1^I1016T^ human fibroblasts and U2OS-SRA-shNPC1 cells expressing *NPC1*^*I1016T*^

WT human fibroblasts (GM05659), NPC1^I1061T/P237S^ (GM03123), and NPC1^I1061T^ (GM18453) human fibroblasts were treated with 10µM Vorinostat or DMSO solvent control for 48 hours. Cells were then incubated with 50 µg/ ml Alexa546-LDL in MEM growth medium supplemented with 5.5% FBS and 10 µM Vorinostat or DMSO solvent control for 4 hours, rinsed with growth medium and chased in MEM growth medium supplemented with 5.5% FBS for 30 minutes. Cells were washed three times with PBS and then fixed with 1.5% PFA in PBS. For immunostaining cells were permeabilized with 0.5% saponin and 10% GS in PBS for 30 minutes. Cells were incubated with 0.8 µg/ml anti-NPC1 rabbit polyclonal primary antibody for two hours in the presence of 0.05% saponin and 0.5% GS at room temperature, followed by Alexa 488 labeled goat anti-rabbit secondary antibody (1:1000, Life Technologies, Grand Island, NY) for 45 min at room temperature. Finally, cells were washed three times with PBS, and images were acquired using a wide-field microscope with a 63X 1.32 NA oil immersion objective and standard FITC and TRITC filters.

U2OS-SRA-shNPC1 cells were transfected with WT or NPC1^I1061T^. One day after transfection, cells were treated with 10 µM Vorinostat, 50 nM Panobinostat or DMSO solvent control. After 48 hours treatment, cells were fixed with 1.5% PFA and processed for immunofluorescence as above except that 180 ng/ml anti-NPC1 rat monoclonal antibody (produced in the Balch laboratory) and 1:500 anti-LAMP1 rabbit polyclonal antibody were used followed by 1:1000 dilution of Cy5 goat anti-rat and 1:500 of Alexa Fluor 546 goat anti-rabbit as secondary antibodies. Fluorescence images were acquired with a Zeiss LSM 510 laser scanning confocal microscope (Thornwood, NY) using a 63X 1.4 NA objective (axial resolution 1.0 μm).

## Results

### Protein lifetime measurements

Treatment with Vorinostat and other HDACi increases the expression of the NPC1^I1061T^ protein in human patient-derived fibroblasts (19). We measured the effect of treatment with Vorinostat on the lifetime of newly synthesized NPC1 protein in fibroblasts with a homozygous NPC1^I1061T^ mutation. As described previously (23), a substantial fraction of the WT NPC1 is degraded rapidly, and the remainder has a half time of over a day (Fig. 1). Nearly all of the NPC1^I1061T^ protein is degraded rapidly in untreated cells. When cells with the NPC1^I1061T^ mutation were pretreated for two days with Vorinostat (10 μM), the protein degradation profile was remarkably similar to that seen for the wild type protein. This indicates that treatment with an HDACi prevents the excessive degradation of the mutant protein.

**Fig. 1:**
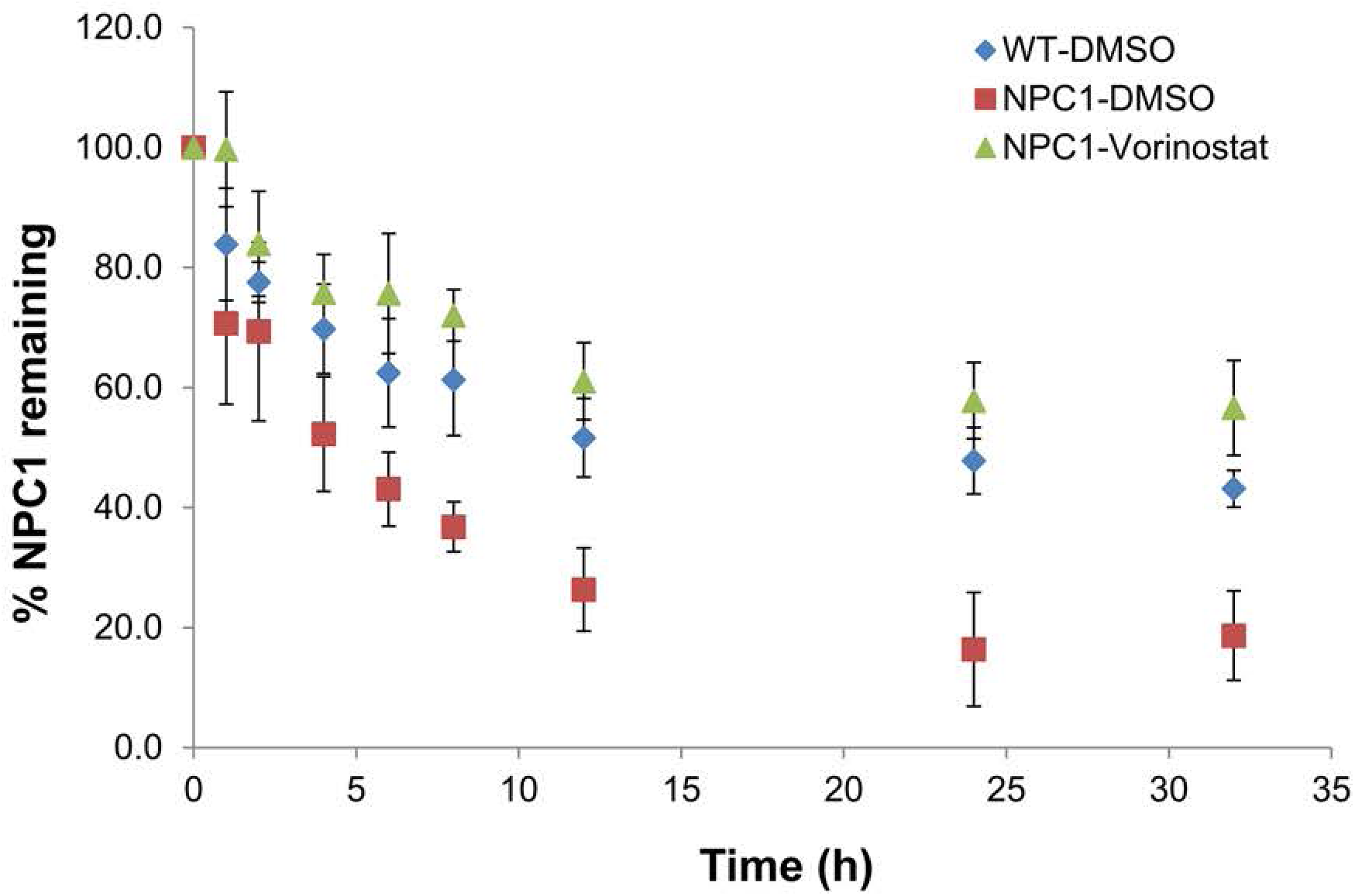
NPC1 protein lifetime measurements. After 48 h treatment with Vorinostat (or DMSO solvent control), GM05659 (WT) or GM18453 (NPC1^I1061T^) fibroblasts were incubated for one hour with culture medium containing [^35^S] Cys/Met followed by a 0–48 h chase in normal culture medium. After the end of each chase time, cells were lysed, and cell lysates were immunoprecipitated with anti-NPC1 antibody. Protein samples were separated by SDS-PAGE, and the densitometry values corresponding to radiolabeled NPC1 were measured. For each experiment the values were normalized to the zero chase time value. Data are shown for solvent-treated GM05659 cells (blue diamonds), solvent-treated GM18453 cells (red squares), and Vorinostat-treated GM18453 cells (green triangles). Each data point is representative of six values from three independent experiments ± SEM.

### HDAC inhibitors lead to correct localization of mutant NPC1

To see if the treatment with HDACi leads to correct targeting of the mutant protein, we examined the localization of mutant NPC1 protein in GM03123 (P237S & splice mutation, I1061T) (Fig. 2A) and in GM18453 (I1061T) (Fig. 2B) *NPC1* mutant human fibroblasts. The cells were treated with 10 μM Vorinostat or solvent control DMSO for 48 hours. The cells were then incubated with Alexa546 LDL for 4 hours to label the LE/Ly that contain lipoproteins. The cells were fixed, and NPC1 protein was detected by immunofluorescence (Fig. 2A and B). Most of the mutant NPC1 protein does not co-localize with LE/Ly in the DMSO-treated cells. After treatment with Vorinostat, a large fraction of the mutant NPC1 protein localizes in LE/Ly.

**Fig. 2:**
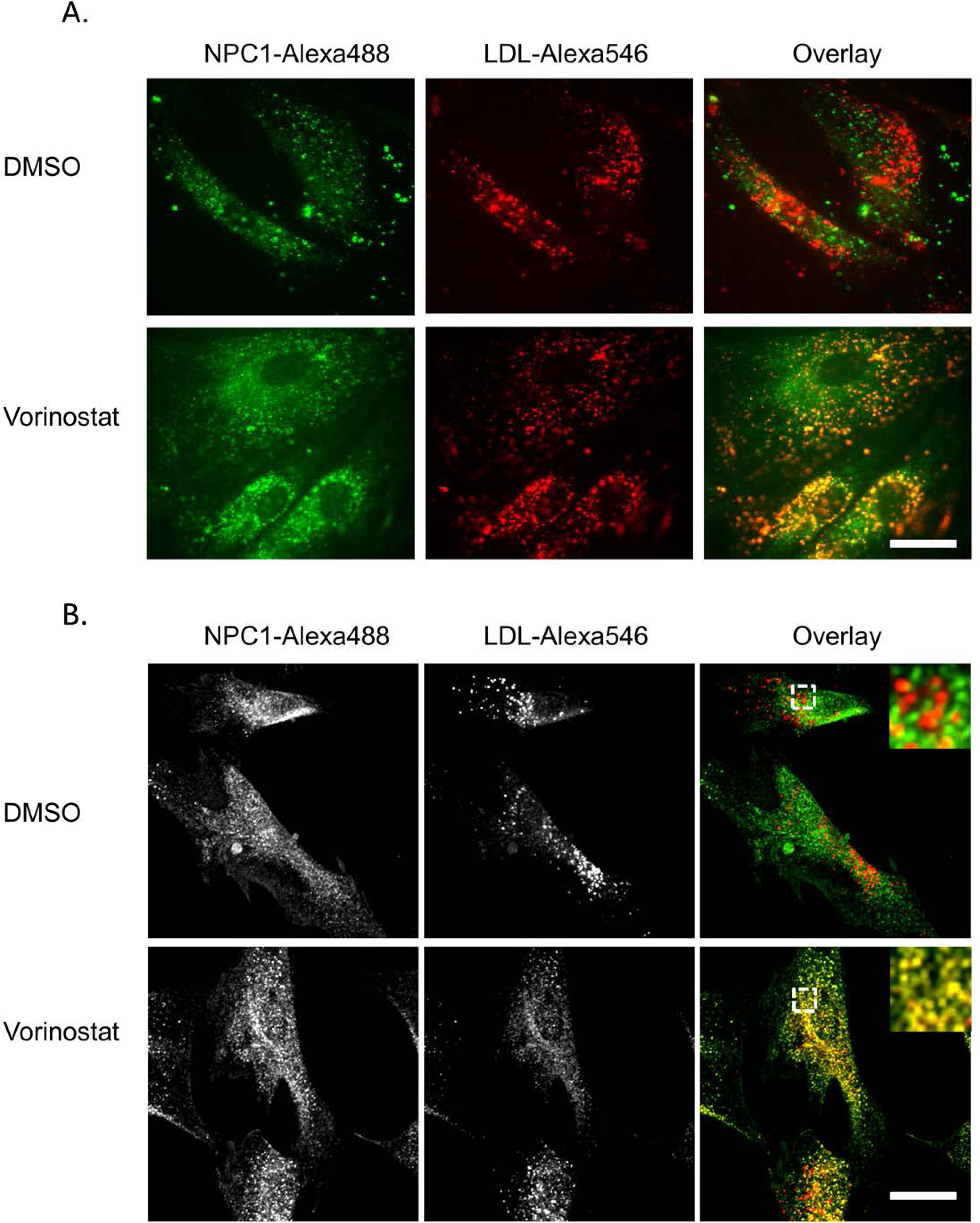
Vorinostat rescues the localization of NPC1 in NPC1 mutant fibroblasts. GM03123 (**A**) or GM18453 (**B**) cells were plated in 2 cm cover-slip bottom PDL coated dishes in MEM growth medium supplemented with 5.5% FBS. On day 2 cells were treated with DMSO or 10μM Vorinostat and incubated for 48 hours. For the last 4 hours, cells were incubated with 50 μg/ml Alexa546 labeled LDL followed by a 30 minute chase in normal medium. Cells were fixed with 2% PFA, permeabilized and immunostained for NPC1. Images were acquired on Zeiss LSM510 confocal microscope using a 63X objective. Images shown are maximum projections. In B the region outlined in the Overlay images is shown at higher magnification in the inset. Scale bar = 10 μm.

### Testing the activity of Vorinostat after drug withdrawal

If the effect of HDACi is to improve delivery of mutant NPC1 protein to LE/Ly, the effect should persist after removal of the drug. To test this, *NPC1* mutant human fibroblasts GM03123 (Fig. 3A) and GM18453 (Fig. 3B) were treated with Vorinostat at varying concentrations for 72 hours. The cells were rinsed and incubated for an additional 0, 1, 2 or 3 days in normal growth medium without Vorinostat. At each time point, cells were fixed with PFA and stained with filipin. We applied a quantitative image analysis method that has been described previously (19, 35) to measure cholesterol storage in the cells. The filipin-labeled cholesterol in LE/Ly is described by a parameter referred to as an “LSO value”, which measures the filipin fluorescence per cell in regions corresponding to the lysosomal storage organelles (see Methods). The LSO values were determined relative to the corresponding DMSO control treatment, and the normalized value was plotted as a function of Vorinostat concentration. Fig. 3A and B shows that the activity of Vorinostat was retained for 2-3 days after withdrawal of drug.

**Fig. 3:**
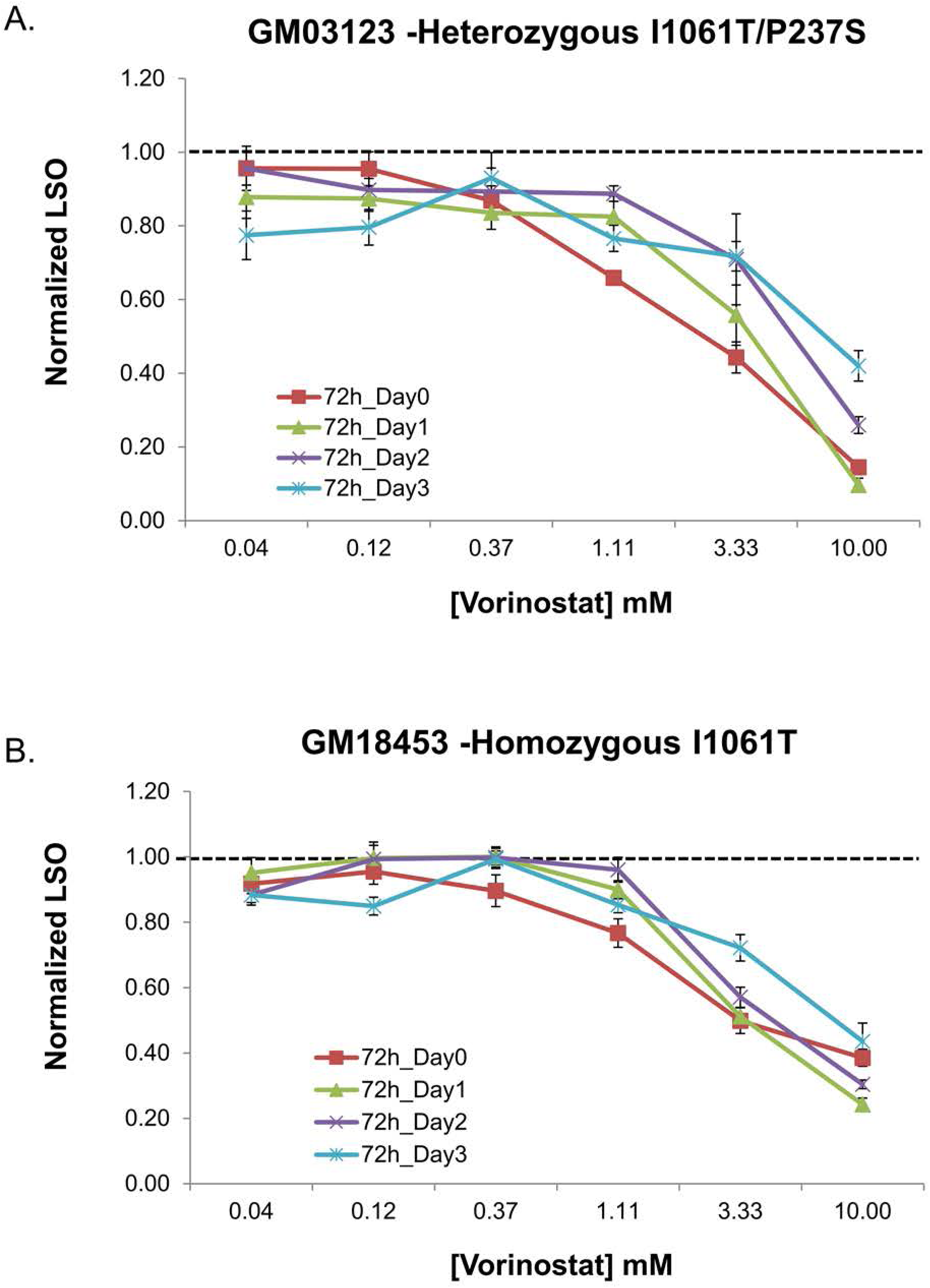
Persistent effect of HDACi treatment in NPC1 mutant human fibroblasts. GM03123 (**A**) or GM18453 (**B**) cells were treated with Vorinostat at varying concentrations for 72 hours. The cells were then incubated for an additional 0, 1, 2 or 3 days without Vorinostat in normal growth medium. At the end of each time point cells were stained with filipin and the LSO value was measured. Data for each cell line are from two independent experiments, and each data point is obtained using 48 images. Each data point is normalized to its corresponding DMSO treated condition, so the value of one represents no effect. Error bars: SEM.

### Treatment of several patient-derived human fibroblasts with Vorinostat

We have shown previously that treatment with several HDACi corrects the cholesterol accumulation in homozygous and heterozygous I1061T mutant cell lines (19). To see if Vorinostat would also correct other NPC1 mutations, we tested its effect on cholesterol accumulation in eight lines of patient-derived fibroblasts (Table 1). All of the cell lines responded to Vorinostat, but the dose response varied among the cell lines (Fig. 4A). Two cell lines (NPC1-17 and NPC1-25) had only partial reduction in cholesterol storage after 48 hours treatment, while the NPC1-22 cells were nearly completely corrected by approximately 1 μM Vorinostat. These results indicate that Vorinostat treatment can correct the NPC1 phenotype in cell lines with several different compound heterozygous mutations. Similar results were obtained with Panobinostat, and NPC1-25 was also only partially responsive to Panobinostat as compared to the other cell lines tested (Fig. 4B). As observed previously (19), high concentrations of Panobinostat lead to higher levels of cholesterol storage.

**Fig. 4.**
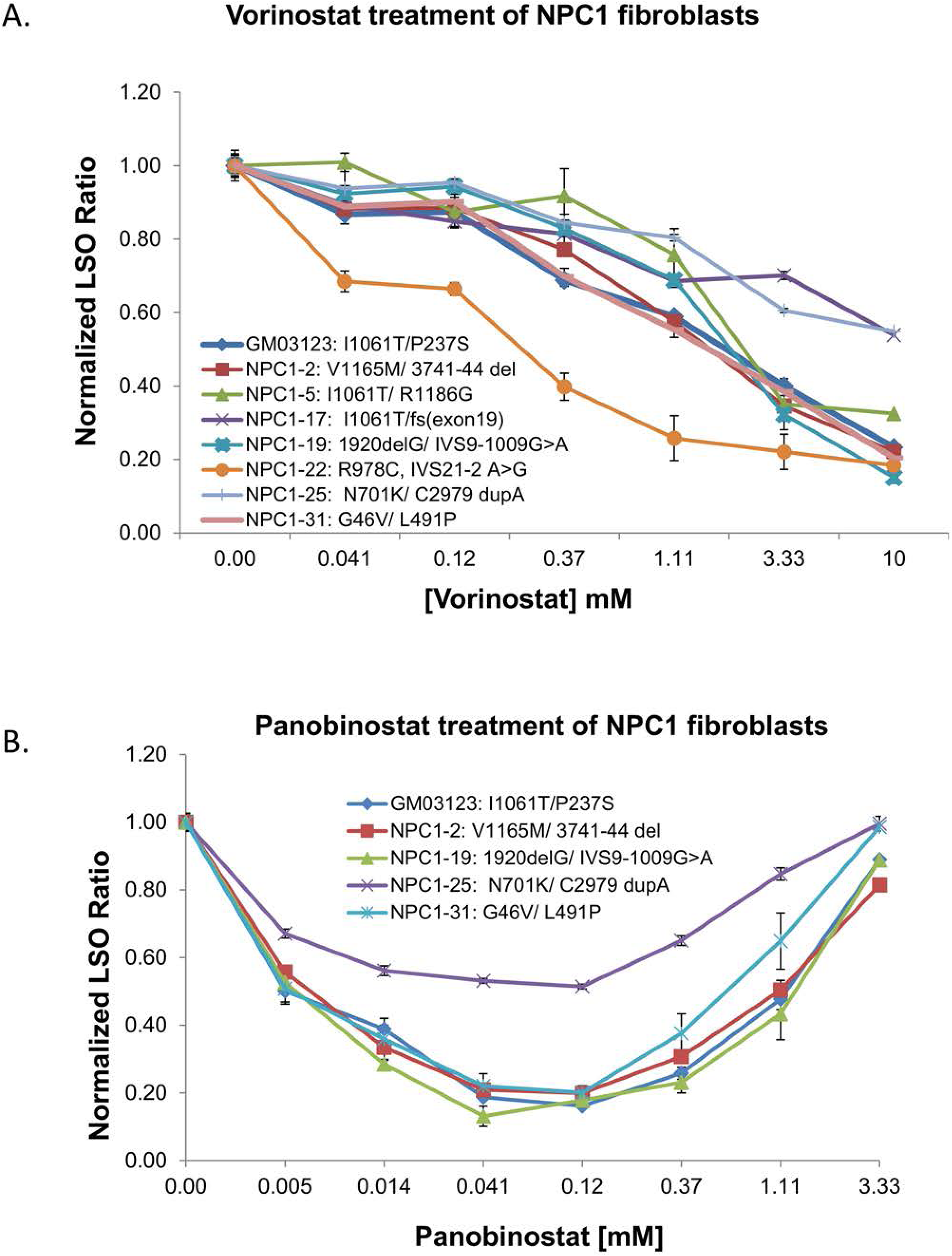
Dose dependent effect of Vorinostat and Panobinostat on multiple patient derived *NPC1* mutant cell lines. NPC1 mutant human fibroblasts were treated with Vorinostat (A) or Panobinostat (B) for 48 hours followed by fixation, staining with filipin and imaging using the ImageXpress^Micro^ automatic fluorescence microscope. DMSO was used as a solvent control. Images were analyzed to obtain the LSO value as a measure of cholesterol accumulation. Data were normalized to the corresponding DMSO treated cells. Data for each cell line are averages of three independent experiments totaling 60 images (5 wells x 4 sites x 3 experiments). Error bars: SEM.

### Effect of HDACi in cells expressing *NPC1* mutants

To test the effect of HDACi on individual NPC1 mutations, we established a cell line that lacked expression of endogenous NPC1 and expressed scavenger receptor type A (SRA) to allow efficient uptake of lipoprotein-derived cholesterol. We stably transfected SRA into a human osteosarcoma cell line, U2OS cells, that have been used widely in drug screening trials (36, 37). These U2OS-SRA cells were then stably transfected with a plasmid expressing shRNA targeting a non-translated region of the NPC1 mRNA to silence the expression of endogenous NPC1. A library of disease-associated *NPC1* mutants was created in a bicistronic vector also expressing GFP to identify the transfected cells. Using reverse transfection, different *NPC1* mutants were transfected into individual wells of a 384-well plate. Half of the wells were treated with an HDACi (in DMSO), while the other half were treated with the same amount of DMSO as a solvent control. Vectors expressing wild type NPC1 and NPC1^I1061T^ were included in each plate as controls.

Fig. 5 illustrates the screening process. After transfection and drug treatment, cells were fixed and stained with filipin, a fluorescent dye that binds to unesterified cholesterol. Untransfected U2OS-SRA-shNPC1 cells incubated with 50 µg/ml AcLDL for 2 hours are brightly labeled with filipin (Fig. 5), reflecting the high levels of unesterified cholesterol stored in the LSOs of these cells (19, 35, 38). Cells transfected with WT NPC1 showed greatly reduced filipin labeling as compared to their untransfected neighbors (Fig. 5). Cells transfected with NPC1^I1061T^ showed little, if any, decrease in filipin labeling as compared to their untransfected neighbors. However, treatment of these NPC1^I1061T^ transfected cells with either Vorinostat (10 µM) or Panobinostat (50 nM) for two days greatly reduced the filipin labeling of transfected cells with no discernible effect on the untransfected neighbors. This indicates that the NPC1^I1061T^ protein can effectively remove cholesterol from LSOs in cells treated with an effective dose of an HDACi.

**Fig. 5.**
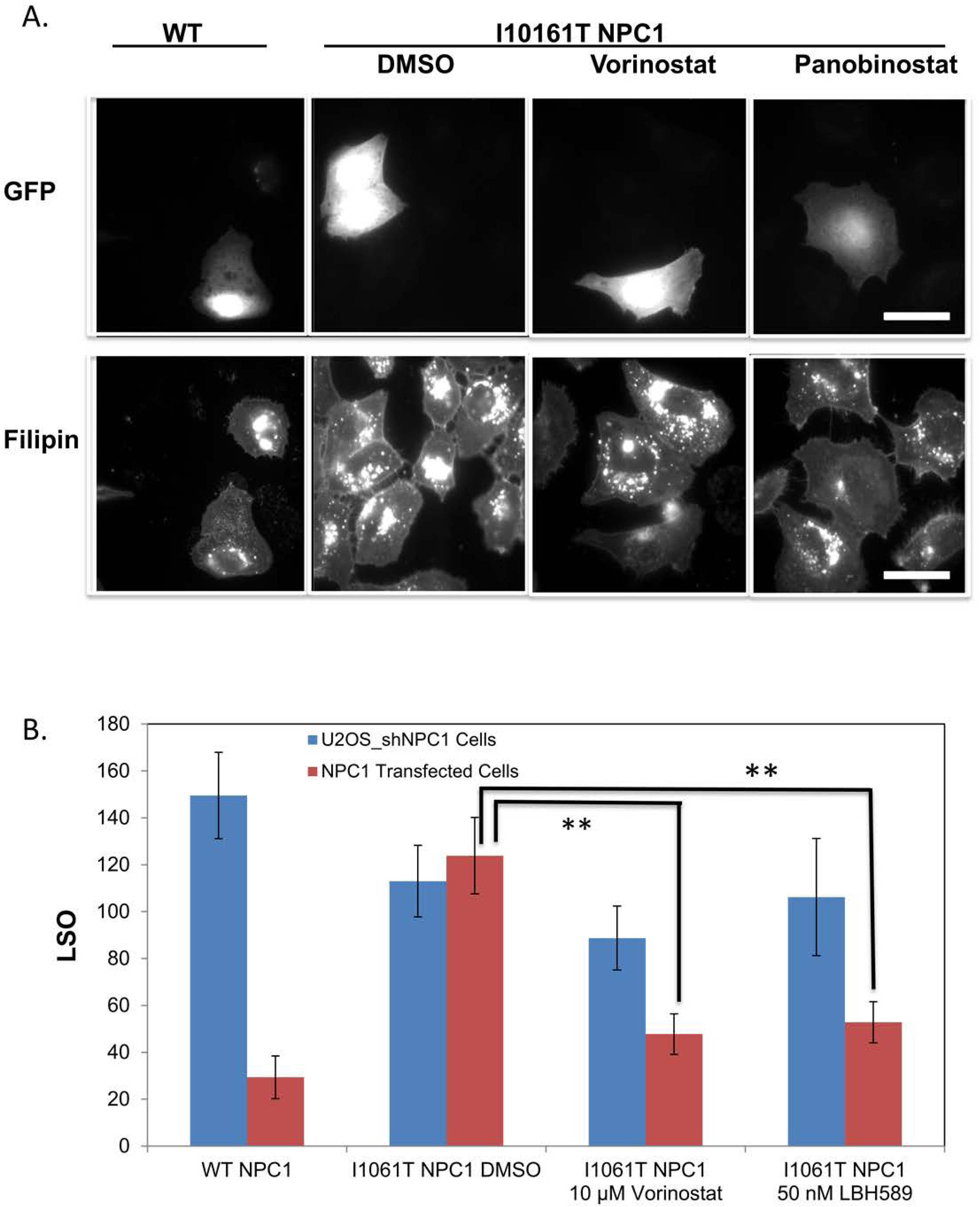
Illustration of the screening system. *A. Representative images of transfected cells.* 24 hours after transfection with a bicistronic vector containing eGFP plus wild type NPC1 or NPC1^I1061T^, U2OS-SRA-shNPC1 cells were treated with Vorinostat (10 µM), Panobinostat (50 nM), or DMSO solvent control for 48 hours. Cells were treated with 50 µg/ml AcLDL for the final 2 hours, fixed with PFA, and stained with filipin. Images were acquired on a Leica wide-field microscope using standard GFP and A4 filters. Transfection with wild type NPC1 reduces the cholesterol accumulation, but transfection with NPC1^I1061T^ does not unless they are treated with an HDACi. *B. Quantification of filipin in transfected cells.* Cholesterol accumulation in LSO of GFP positive cells was measured based on filipin intensity, and the LSO values are shown. Error bars: SEM.

To quantify the relative amount of cholesterol storage under various conditions, we used an automated image analysis protocol based on filipin imaging as described previously (19, 35). As described in Methods, an “LSO value” is determined, which reflects the amount of filipin observed in brightly labeled regions of the cells corresponding to LSOs. Fig. 5B shows the quantitative analysis of the effects of transfecting U2OS-SRA-shNPC1 with wild type NPC1 or NPC1^I1061T^. Expression of the NPC1^I1061T^ mutant only caused a significant decrease in the LSO value in cells treated with an HDACi. As with the GM03123 human patient fibroblasts, treatment of U2OS-SRA-shNPC1 cells expressing NPC1^I1061T^ with an HDACi leads to delivery of the mutant NPC1 to LE/Ly labeled with LAMP1 (Fig. 6).

**Fig. 6.**
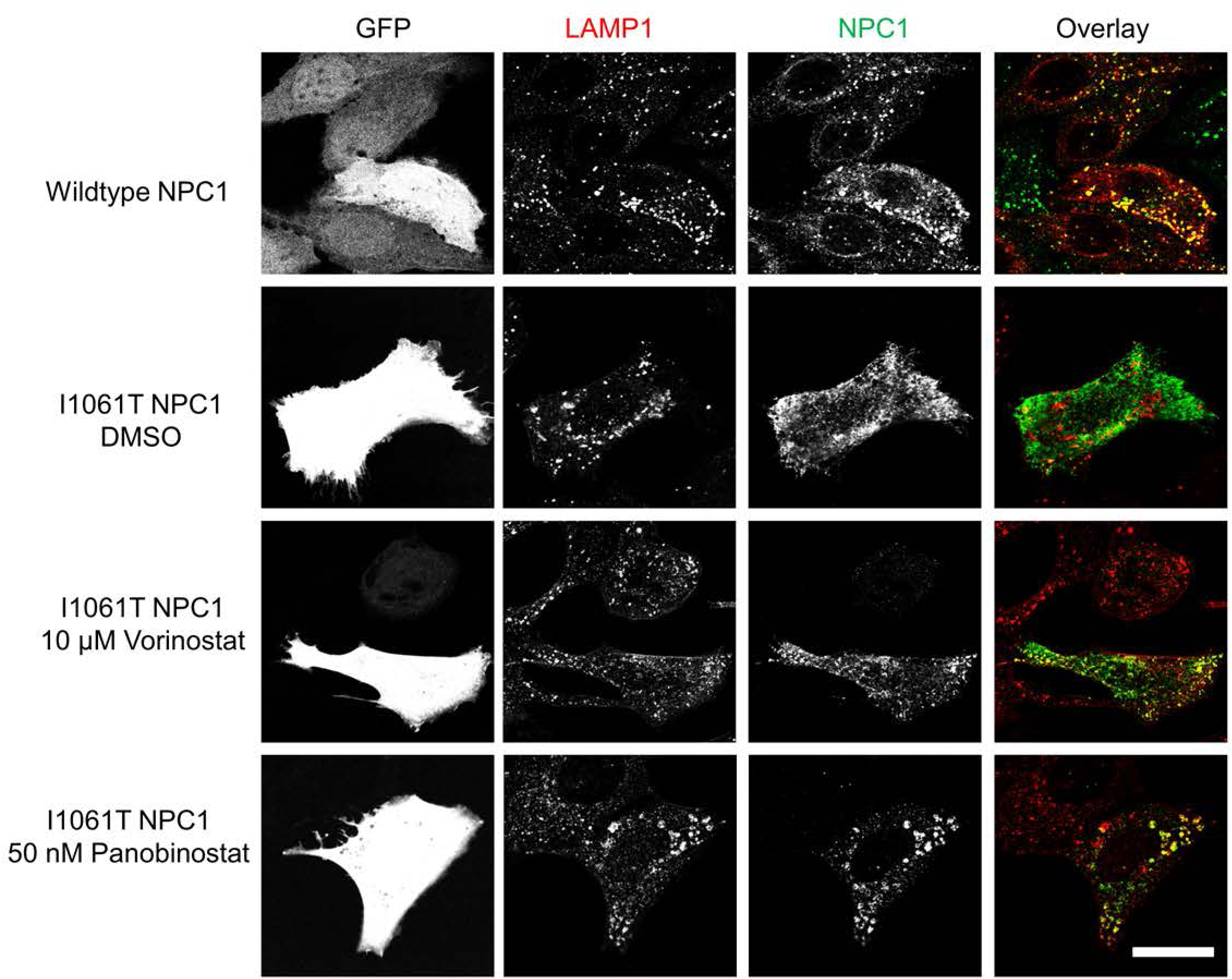
Vorinostat and Panobinostat rescue localization of NPC1^I1061T^ in U2OS-SRA-shNPC1 cells. WT NPC1 or NPC1^I1061T^ was expressed in U2OS-SRA-shNPC1 cells using a bicistronic vector also encoding eGFP. Starting one day later, wells transfected with NPC1^I1061T^ were treated with 10 µM Vorinostat, 50 nM Panobinostat or DMSO solvent control for 48 hours. GFP serves as a marker of transfected cells. NPC1 protein was detected by immunofluorescence using anti-NPC1 Rat monoclonal primary antibody and Cy5 Goat anti-rat secondary antibody. Lysosomes were identified by using anti-LAMP1 rabbit polyclonal antibody and Alexa Fluor 546 goat anti-rabbit secondary antibody. The co-localizations of NPC1 protein and LE/Ly are shown in yellow. Scale bar = 10 μm

Using the assay for filipin labeling, we tested the effect of Vorinostat (10 μM) and Panobinostat (50 nM) on 81 different NPC1 mutations that have been found in patients (Fig. 7 and **Supplementary Fig. S1**). With Vorinostat treatment, 68 of the 81 mutants showed a reduction of the LSO Value with p < 0.05. The results are summarized in Fig. 8. Mutants that showed a statistically significant (p < 0.05) response to Vorinostat are listed in black, and nonresponsive mutants are listed in red. Over 80% of the mutations showed a statistically significant response to either of the HDACi treatments. Some of the mutants that were not corrected to a statistically significant level had low starting cholesterol, which made it more difficult to demonstrate an effect. Expression of these mutants may have had some corrective effect even before HDACi treatment. In general the data shown in Fig. 7 indicate that there is not a strong correlation between the LSO value before treatment and the correction seen after HDACi treatment. After HDACi treatment, many of the mutants that showed high cholesterol storage before treatment were corrected to nearly the level in WT transfectants.

**Fig. 7.**
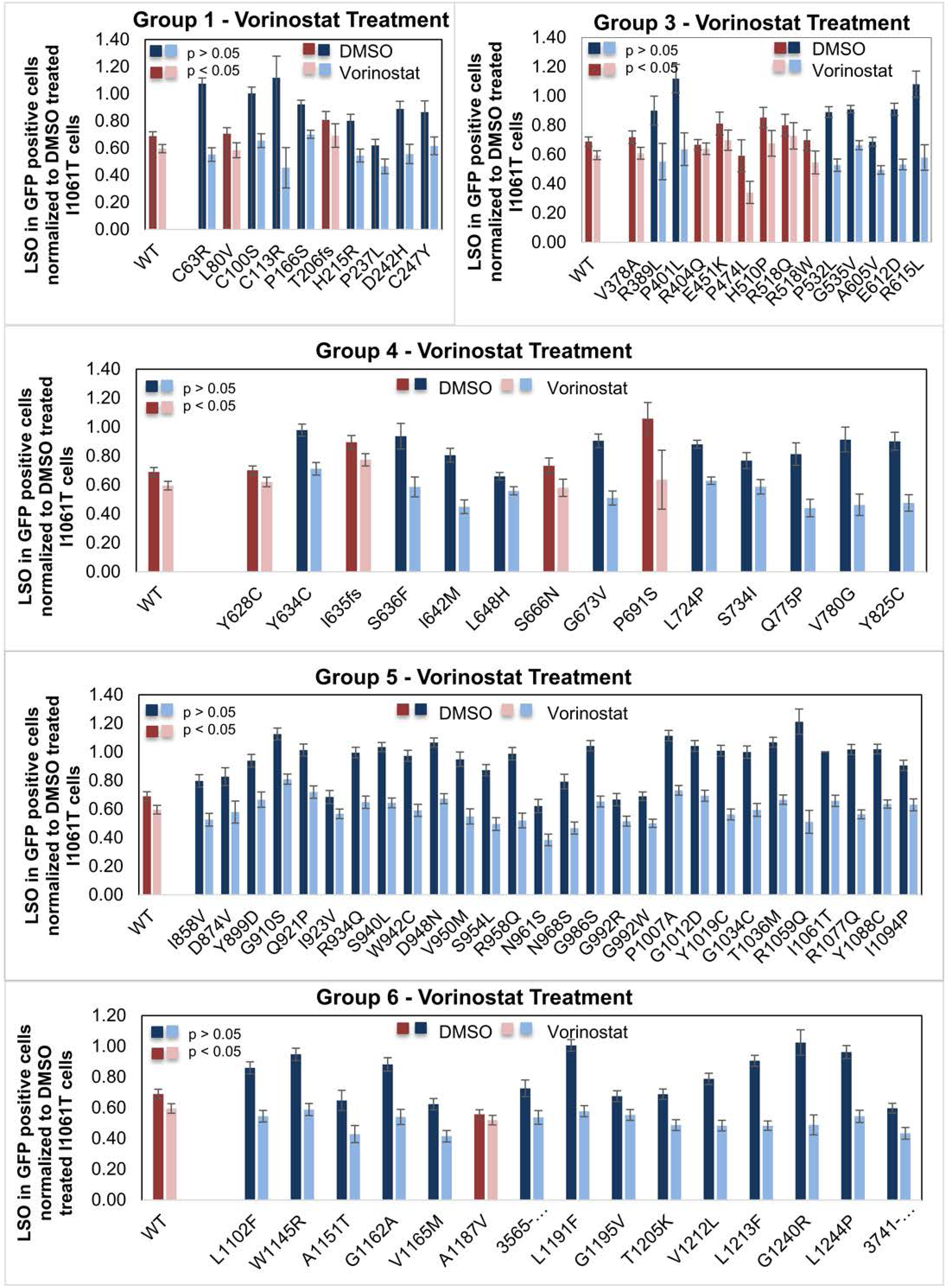
Quantification of effects of Vorinostat on cholesterol accumulation in NPC1 mutants. The effect of 10 μM Vorinostat was tested on 81 different NPC1 mutations from five segments of the NPC1 protein as illustrated in Figure 8. DMSO was used as a solvent control. Filipin fluorescence images of the transfected cells were analyzed to obtain an LSO value as explained in methods. Data represent averages ± SEM from 15-25 images. Each image includes about 1-5 transfected cells. Dark blue (DMSO treated) and light blue bars (Vorinostat treated) for each group represent mutants that showed reduction in LSO values with p<0.05, and dark red (DMSO treated) and pink bars (Vorinostat treated) represent mutants that showed no significant reduction in LSO value. Statistical significance was measured by t-test with two-tailed distribution and two-sample equal variance (homoscedastic) type using Microsoft Excel software.

**Figure 8.**
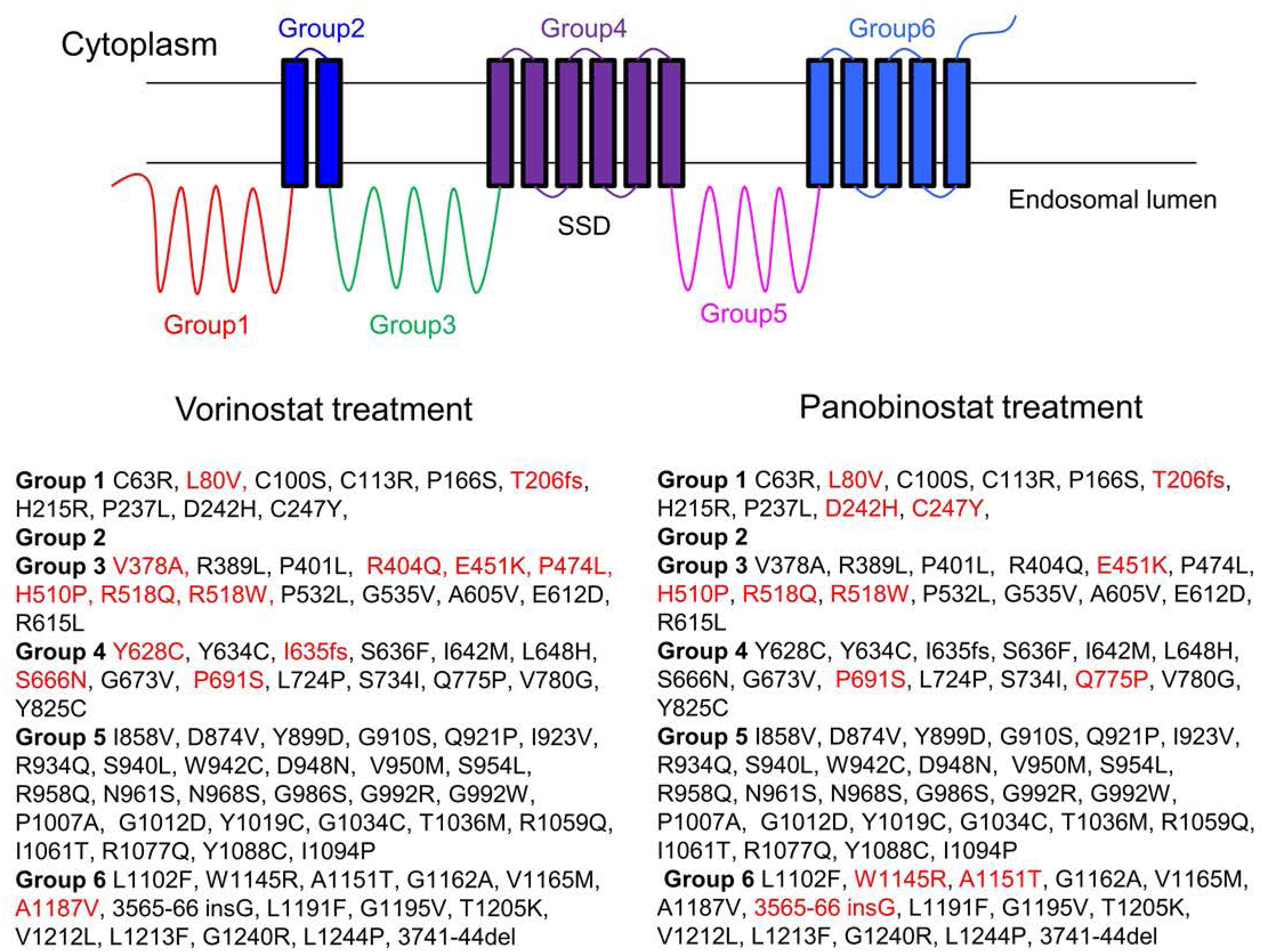
List of NPC1 mutants tested with Vorinostat and Panobinostat. 81 NPC1 mutants were treated with Vorinostat (10 μM) or Panobinostat (50 nM) as described in Fig. 7. The mutants are separated into 6 groups by their locations in the NPC1 protein structure. Mutants that showed a statistically significant (p < 0.05) response to Vorinostat are listed in black, and nonresponsive mutants (p > 0.05) are in red.

A full listing of the mutants and the statistical significance of their responses is provided in **Supplementary Table 1**. To see if correction depended on the severity of the initial cholesterol accumulation, we compared the LSO values of DMSO-treated and Vorinostat or Panobinostat treated cells. It can be seen that most of the mutations with the highest initial cholesterol storage were corrected.

## Discussion

Niemann-Pick C disease is a devastating inherited disorder with no US FDA approved treatment. More than 300 disease causing mutations have been found in NPC1 (10, 30), and most of these are missense mutations in this large membrane protein. As part of a screen for compounds that would correct the cholesterol storage defect in patient-derived fibroblasts, we found that several HDACi compounds were included among the hits (19). The pharmacological profile of the effective inhibitors suggested that HDACs 1, 2, or 3 were the most relevant targets (19, 20). Although increased expression of the NPC1^I1061T^ protein was observed following treatment with HDACi, the mechanism of correction and the applicability to a variety of mutations were not determined.

It has been shown previously that nearly half of the wild type NPC1 protein is degraded by an ERAD pathway, and essentially all of the newly synthesized NPC1^I1061T^ protein is degraded by ERAD (23). In this study we found that treatment of cells with Vorinostat causes the stability of the NPC1^I1061T^ protein to be indistinguishable from the wild type protein. The longer-lived fraction of both wild type and HDACi-treated NPC1^I1061T^ are degraded with a halftime of over one day. We also observed that the NPC1^I1061T^ protein is correctly delivered to LE/Ly that contain endocytosed LDL, so after exit from the ER a significant amount of the mutant protein is trafficked to the correct organelle. The effect of HDACi treatment on cholesterol storage in LSOs persists for 2-3 days after withdrawal of the drug, which is consistent with the long lifetime of the NPC1 protein after it has passed the ER quality control system. The long term effect of the HDACi after removal from the medium may be useful in designing clinical protocols for treatment.

In order to test the effect of HDACi treatment on a large number of mutants, we used a reverse transfection strategy that could be used for screening in multiwell plates (39). The U2OS-SRA-shNPC1 cells were developed for this purpose. The stable expression of SRA was valuable since it allowed us to deliver a large amount of cholesterol to the cells via AcLDL, which binds to SRA and is internalized by receptor-mediated endocytosis (40). Cells with defects in the NPC1 protein were unable to clear this bolus of cholesterol from the LE/Ly. The bicistronic vector containing eGFP and NPC1 allowed us to identify cells that were transfected with the wild type or mutant NPC1. The reverse transfection method we used gave about 3% transfection efficiency on the average, which allowed us to observe transfected and untransfected cells in the same microscope fields.

It was shown previously that very high levels of over-expression of the NPC1^I1061T^ protein can correct the cholesterol storage defect in cells, presumably because a small fraction of the protein escapes ERAD and gets onto the LE/Ly (23). We observed that more than 80% of the NPC1 mutants could be corrected for their cholesterol accumulation by treatment with either Vorinostat or Panobinostat. This would indicate that more than 90% of patients would be expected to have at least one allele that is susceptible to HDACi treatment. Mechanistically, this suggests that the large majority of NPC1 mis-sense mutations can be functional if they are delivered to the correct organelles.

The analysis of the mutants that can and cannot be corrected provides some insight into the requirements for a functional NPC1 protein. One caveat is that these assays were carried out in a high throughput screening system, and they have not been individually verified in detail. However, analysis of the data provides some measures of quality control. First, it should be noted that there is a good general correspondence between the mutations that were affected by Vorinostat and those affected by Panobinostat. Some of the mutants that were listed as corrected by one drug but not the other were only statistically significant at p < 0.05 (Supplementary Table 1), suggesting that these might have been only partially corrected by one or both HDACi. The frame shift that occurs in the position corresponding to T206 results in a stop codon and is effectively an NPC1 null. The lack of effect of both Vorinostat and Panobinostat is consistent with the lack of an effect of HDACi treatment on cells lacking NPC1 expression. One of the non-corrected mutations (L80V) was in the N-terminal domain, which is responsible for cholesterol binding. The leucine at position 80 is close to the cholesterol binding pocket (4). NPC2 binds to the first lumenal loop (Group 3 in Fig. 8) of NPC1, and it has been shown that mutations R404Q and R518Q interfere with this binding (41). Neither of these mutations was corrected by Vorinostat, and R518Q is not corrected by Panobinostat. It has been shown that R518Q goes to punctate organelles in a distribution consistent with LE/Ly (41). The P691S mutation is in the sterol sensing domain of NPC1 (28). It has been shown that this mutation is ineffective in promoting cholesterol transport even though it is delivered correctly to LE/Ly (42). It is noteworthy that all of the mutations in Group 5 can be rescued by both Vorinostat and Panobinostat. Apparently this segment of NPC1 can tolerate many mutations without losing the ability to export cholesterol from LE/Ly.

**Table 1:** List of mutations in NPC1 patient fibroblasts

|  |  |
| --- | --- |
| GM03123 | Heterozygous I1061T/ P237S |
| GM18453 | Homozygous I1061T |
| NPC1-2 | V1165M/ 3741-44 del ACTC |
| NPC1-5 | I1061T/ R1186G |
| NPC1-16 | P887L/ 3741-44 del ACTC |
| NPC1-17 | I1061T, 10 bp deletion in exon 19 at codon 962=fs(exon19) |
| NPC1-19 | 1920delG/ IVS9-1009G>A |
| NPC1-22 | R978C, IVS21-2 A>G |
| NPC1-25 | N701K/ C2979 dupA |
| NPC1-31 | G46V/ L491P |

The mechanism by which HDACi treatment rescues the function of mutant NPC1 proteins will require further mechanistic studies. One possibility is that HDACi treatment alters the proteostatic environment. It has been reported that HDACi treatment can increase expression of some protein chaperonins (43) or directly modulate the activity of chaperones by altering their acetylation status (44).

An indirect mechanism for overcoming the cholesterol accumulation caused by mutations in *NPC1* is consistent with findings that patients with identical mutations can have different times to initial onset and disease severity (10). This suggests that other factors in the genetic background can alter the degree of loss of function. It is not known if this is related to higher levels of correct targeting of mutant NPC1 protein in the tissues of less affected individuals.

Our data suggest that the vast majority of NPC1 patients carry mutations that could be corrected by HDACi treatment. Vorinostat and Panobinostat are FDA-approved drugs for treatment of some cancers, and HDACi have been used in clinical trials for many other diseases (20). Vorinostat is generally well tolerated as a cancer therapeutic agent (45). Several other HDACi have been in large scale clinical trials. Since the most important pathologies of NPC disease are related to neuronal cell dysfunction and death (10), penetration of an HDACi into the brain will be essential for effective treatment. Some HDACi have excellent penetration into the brain and have been used in animal and human studies for treatment of neurological diseases (20). The results described herein support the investigation of HDACi as single or combination therapies (27) for NPC1 disease.

## Acknowledgements

We thank Dr. Forbes D. Porter (NICHD, Bethesda, MD) for supplying several NPC patient fibroblast cell lines, and we also thank Dr. Dan Ory (Washington University, St. Louis, MO) for providing NPC1 cDNA. We thank Dr. Paul Helquist (University of Notre Dame, South Bend, IN) for supplying Vorinostat and Panobinostat. This work was supported by grant R01-NS092653 from the NIH and grants from the Ara Parseghian Medical Research Fund to FRM.

